# Microaerophilic activated sludge system for ammonia recovery from high-strength nitrogenous wastewater: Performance and microbial communities

**DOI:** 10.1101/2022.07.21.500714

**Authors:** Hiroki Tsukamoto, Hop Van Phan, Toshikazu Suenaga, Shohei Yasuda, Megumi Kuroiwa, Shohei Riya, Atsushi Ogata, Tomoyuki Hori, Akihiko Terada

## Abstract

A transition to ammonia recovery from wastewater has started; however, a technology for sustainable nitrogen retention in the form of ammonia is still in development. This study validated a microaerophilic activated sludge (MAS) system to efficiently retain ammonia from high-strength nitrogenous wastewater. The MAS is based on conventional activated sludge (CAS) with aerobic and settling compartments. Low dissolved oxygen (DO) concentrations (<0.1 mg/L) and short solid retention times (SRTs) (<5 d) eliminated nitrifying bacteria. The two parallel MASs were successfully operated for 300 d and had ammonia retention of 101.7 ± 24.9% and organic carbon removal of 85.5 ± 8.9%. The MASs mitigated N_2_O emissions with an emission factor of <0.23%, much lower than the default value of CAS (1.6%). A short-term step-change test demonstrated that N_2_O indicated the initiation of nitrification and the completion of denitrification in the MAS. The parallel MASs had comparable microbial diversity, promoting organic carbon oxidation while inhibiting ammonia-oxidizing microorganisms (AOMs), as revealed by 16S rRNA gene amplicon sequencing, qPCR of functional genes, and fluorescent *in situ* hybridization of β-Proteobacteria AOB. The microbial analyses also uncovered that filamentous bacteria were positively correlated with effluent turbidity. Together, controlling DO and SRT achieved successful ammonia retention, mainly by suppressing AOM activity. This process represents a new nitrogen management paradigm.

**Synopsis:** Moving from nitrogen removal to nitrogen recovery is critical for establishing a sustainable society. We provided proof-of-the-concept for a novel ammonia retention technology by retrofitting an activated sludge system.

## Introduction

Human activities have converted a massive amount of nitrogen gas into reactive nitrogen (Nr) via the Haber−Bosch process for food production in response to the surging world population.^1^ Approximately 50% of Nr remains in the environment, causing pollution in water systems, coastal areas, and soils.^2^ The accumulated volume of Nr in the environment has exceeded what the natural environment can accommodate and cannot be reversed.^3^ Given that the Haber−Bosch process is energy-intensive, requiring as much as 43.7 MJ/kg-N^4^ (equivalent to 1.6% of the total CO_2_ emissions^5^), Nr production by this process should be reduced while not compromising anthropogenic activities. To ensure the global food supply, the recovery and reuse of Nr compounds from waste streams is indispensable.

A sustainable solution for nitrogen management in water industries is urgently needed. An estimated 20 Mt/year of ammonia is released in municipal wastewater, equivalent to 19% of the Nr compounds anthropogenically fixed by the Haber−Bosch process.^6^ Ammonia is conventionally removed during wastewater treatment to prevent eutrophication in receiving water bodies. Biological ammonium removal, which converts ammonia into inert nitrogen gas mainly via nitrification and denitrification, is energy intensive. Aeration required by nitrification accounts for up to 50% (45 MJ/kg-N) of the operational cost of wastewater treatment plants (WWTPs),^7^ and is a comparable energy consumption to ammonia production via the Haber−Bosch process.^8^ Therefore, shifting ammonia removal from wastewater to ammonia recovery is essential for the sustainable management of Nr compounds.^9^

Some ammonia recovery technologies have been developed and implemented, including ammonium stripping, struvite precipitation, bio-electrochemical systems,^10^ membrane separation,^11^ ion exchange, and adsorption.^12^ These technologies allow the recovery and reuse of Nr compounds, but still require substantial energy and/or chemical resources, which increases operational costs and environmental burdens.^13^ A sustainable solution that allows ammonia recovery from wastewater using less energy has yet to be developed. Furthermore, of importance is, prior to these ammonia recovery technologies, retaining ammonia in wastewater that contains multiple nitrogen constituents. Therefore, this study proposes a novel process, termed a microaerophilic activated sludge (MAS) system, for removal of organic compounds and retention of ammonia. The principle of this technology is to enrich ammonia by suppressing ammonia oxidation and by converting organic nitrogen compounds to ammonia. The MAS system, where ammonia is recovered in a downstream process, can save energy by avoiding conventional ammonia removal and by retrofitting WWTP configurations.

The most important feature of the MAS is inhibiting ammonia oxidation while oxidizing organic carbon. Complex and functionally diverse microbial guilds are present in activated sludge, and ammonia oxidation via nitrite to nitrate, termed nitrification, is carried out by nitrifying microorganisms. These include canonical ammonia-oxidizing bacteria and archaea (AOB and AOA), complete ammonia-oxidizing (commamox) bacteria belonging to *Nitrospira*, and nitrite-oxidizing bacteria (NOB). The MAS system eliminates these ammonia-oxidizing microorganisms (AOMs) based on their physiological properties. AOMs have much lower specific growth rates (*µ*_*max*_ [h^−1^]) (0.028–0.085 h^−1^ for AOB; 0.010– 0.013 h^−1^ for AOA; 0.019–0.060 h^−1^ for NOB, and 0.006 h^−1^ for comammox *Nitrospira*)^14^ and oxygen affinities (oxygen half-saturation coefficients *K*_*O*_ [mM]) (1–40 mM for AOB, 2–4 mM for AOA, and 4.06–16.88 mM for NOB)^14^ than those of heterotrophic bacteria (typical *µ*_*max*_ of 0.25 h^−1^ and *K*_*O*_ of 0.00312 mM).^15, 16^ These distinctive biokinetic properties should allow the suppression of AOMs growth but sustain heterotrophic bacteria growth by controlling operational parameters (*e*.*g*., shorter solid retention time [SRT] and lower dissolved oxygen [DO] concentration^17^). However, this hypothesis requires experimental verification.

Mitigating N_2_O emissions is another critical issue for sustainable nitrogen management because N_2_O, a highly potent greenhouse and ozone-depleting substance,^18^ is produced as a byproduct and intermediate during nitrification and denitrification, respectively. WWTPs account for 2.8% of total anthropogenic N_2_O emissions,^19^ with an average emission factor of 1.6%.^20^ An N_2_O emission factor of 0.2% from an aeration tank at WWTP is equivalent to approximately 8% of the tank’s CO_2_ footprint.^21^ Although suppressing nitrification theoretically prevents N_2_O formation, unwanted nitrification and N_2_O emissions may still occur under certain conditions. Previous studies reported high N_2_O emissions at a DO concentration of 0.8 mg/L for nitrification and 0.4 mg/L for denitrification.^22^ To efficiently retain ammonia while minimizing N_2_O emissions, a systematic investigation is needed to better understand the N_2_O emission dynamics under low DO conditions. Given that N_2_O emissions were previously reported as an early warning for decreased nitrification efficiency,^23^ understanding the N_2_O emission patterns and the underlying mechanisms can aid in the development of a monitoring tool to prevent unexpected nitrification.

We present the proof-of-concept for a MAS system that can efficiently retain ammonia from high-strength nitrogenous wastewater. The MAS system was operated and established by adjusting the SRT and DO concentrations. Online monitoring of NH_4_^+^, NO_3_^−^, N_2_O, DO, and oxidation-reduction potential (ORP) was implemented. The microbial community and spatial distribution of AOB in activated sludge were analyzed by 16S rRNA gene amplicon sequencing, quantitative PCR (qPCR), and fluorescent *in situ* hybridization (FISH), which confirmed the suppression of AOM growth. A short-term step-change in aeration rate was introduced to investigate if gaseous N_2_O could be used as an indicator for the initiation of nitrification. Together, this study demonstrated the feasibility of the MAS system for highly efficient ammonia retention while minimizing N_2_O emissions, and it revealed the underlying mechanism of the new ammonia retention technology.

## Materials and Methods

### Bioreactor design, setup, and continuous operation

Two laboratory-scale continuously mixed MAS reactors (designated R1 and R2), each with a working volume of 10 L, were constructed and operated. Each reactor consisted of two compartments for aeration (7.2 L) and sedimentation (2.8 L) (**Fig. S1**). The aeration compartment continuously received synthetic fermentation industry wastewater consisting of (per L tap water): 50 mg KH_2_PO_4_, 3.2 mg MgSO_4_·7H_2_O, 20 mg KCl, 200 mg NaHCO_3_, 10 mg NaCl, 1.834 g NH_4_Cl, 0.8 mg FeCl_3_·6H_2_O, 1.5 mg CaCl_2_, 0.5 g tryptone, and 0.5 g yeast extract. The wastewater (before addition of NaHCO_3_) was autoclaved, followed by the addition of NaHCO_3_, before feeding to the system. The two reactors were operated in parallel for approximately 300 d. Each reactor was seeded with activated sludge biomass from a municipal WWTP (Kinu Aqua Station, Ibaraki, Japan) and the synthetic wastewater was fed at a hydraulic retention time (HRT) of 11.2 h. The median influent carbon loading was 0.42 kg-C/m^3^/day. Aeration rate was consistently maintained at 2.0 L/min throughout the operation, except for the period when the short-term step-change experiment was performed. The startup period lasted for 25 d to ensure a mixed liquor suspended solids (MLSS) concentration of above 1000 mg/L. Subsequently, the aeration rate was adjusted by a mass flow meter to maintain a DO < 0.2 mg/L. SRT was set to <5 d by manually withdrawing biomass from the aeration compartment. MLSS was measured twice a week to determine the amount of sludge withdrawal from the aeration compartment.

### N_2_O monitoring by perturbating oxygen loadings

We hypothesized that the rapid changes in exhausted gaseous N_2_O indicated the start point of nitrification and the endpoint of denitrification. To verify this hypothesis, a stepwise change in DO concentration was employed for R1 on day 58, and the corresponding changes in the concentrations of gaseous N_2_O were recorded online. The NH_4_^+^, NO_2_^−^, and NO_3_^−^ concentrations before the stepwise change were 330.3, 2.4, and 16.6 mg-N/L, respectively, which allowed both nitrification and denitrification to be affected by the changing DO concentrations. The details of the experimental procedure are presented in **Table 1**. Throughout the experiment, the dynamic responses of all the important parameters (ORP, pH, DO, and exhausted gaseous N_2_O) were monitored to determine the factors for the activation of nitrification at Phase 4 and the completion of denitrification at Phase 2.

**Table 1:** The experimental procedures of the stepwise change in DO concentration.

| Experimental phases | Objectives | Aeration rate [L/min] | Duration [h] |
| --- | --- | --- | --- |
| Phase 1 | Oxic phase for nitrification | 2.0 | 1 |
| Phase 2 | Anoxic phase for denitrification | 0.5 | 5 |
| Phase 3 | Extended anoxic phase | 0.5 | 2 |
| Phase 4 | Oxic phase for nitrification | 2.0 | 12 |

### Chemical analysis

Samples were collected twice a week to monitor the performance of the system. Total organic carbon (TOC) was measured in samples filtered through glass filters (1825-025, Sigma-Aldrich, Maidstone, UK); disposable membrane filters (25AS045AN, Advantec, Tokyo, Japan) were used to filter samples for NH_4_^+^, NO_2_^−^, and NO_3_^−^ analysis. Dissolved TOC and total nitrogen (TN) concentrations were measured using a TOC analyzer with a TN unit (TOC-L_CPH/CPN_, Shimadzu, Kyoto, Japan). NH_4_^+^, NO_2_^−^, and NO_3_^−^ concentrations were measured by ion chromatography (ICS1000 and Aquion, Thermo Fisher Scientific, Sunnyvale, CA). The exhausted gas in the aeration compartment was collected with an airtight syringe to measure gaseous N_2_O concentration using a quadrupole gas chromatography–mass spectrometry (GCMS-QP2010 Ultra, Shimadzu, Kyoto, Japan). Other parameters (pH, DO, ORP, and temperature) were monitored online by dedicated sensors (YELS-01PH, YELS-01DOY, and ELS-01OR Yamagata Toa DKK, Yamagata, Japan). Turbidity of the effluent was measured with a turbidity meter TBD700 (AS ONE Corporation, Japan). MLSS and mixed liquor volatile suspended solids (MLVSS) were measured according to the Standard Methods.^24^

For the stepwise change experiment, NH_4_^+^ and NO_3_^−^ concentrations were recorded with online electrodes (VARiON plus 700 IQ, Xylem, Washington DC, USA) on day 56. Exhausted gaseous N_2_O was measured online with quantum cascade–laser absorption spectrometry (QC-LAS) (ABB, Zurich, Switzerland).

### Respirometric assay for nitrification

The respirometry assay for nitrification activity in the suspended biomass was conducted using a method previously described.^25^ Biomass samples (30 mL) were collected and washed with 0.02 × phosphate-buffered saline (PBS), followed by adding oxygen-saturated 0.02 × PBS, trace element solution, and NaHCO_3_ of 8330 mg/L. The compositions of the trace element solution (per L) were: 2.2 mg FeSO_4_·7H_2_O, 1.1 mg ZnSO_4_·7H_2_O, 0.53 mg CuCl_2_·2H_2_O, 0.20 mg NiSO_4_·6H_2_O, and 2.2 mg MnSO_4_·6H_2_O. A concentrated NH_4_^+^ solution (1000 mg-N/L) was added to the mixture to a final concentration of 100 mg-N/L. The DO concentration was monitored using a DO electrode (YELS-01DOY). The linear regression of DO concentration versus time and MLVSS concentration was used to calculate a biomass-specific oxygen consumption rate representing the nitrification activity of the biomass.

### TOC removal efficiency, NH_4_^+^ retention efficiency, and N_2_O emission factor

TOC removal and NH_4_^+^ retention efficiencies (%) were defined as Eq. 1 and 2, respectively.

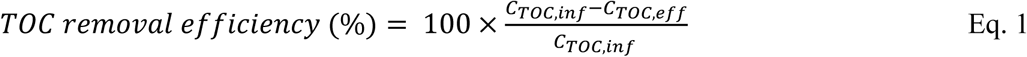

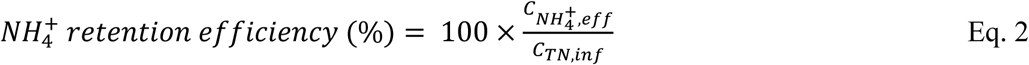

*C*_*TOC,inf*_ (mg-C/L) = Influent dissolved TOC concentration

*C*_*TOC,eff*_ (mg-C/L) = Effluent dissolved TOC concentration

*C*_*TN,inf*_ (mg-C/L) = Influent dissolved TN concentration

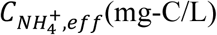 = Effluent NH_4_^+^ concentration

The N_2_O emission factor (%), defined in previous work,^26^ was used with a slight modification (Eq. 3). The factor was the ratio of the emitted rate of the exhausted gas N_2_O and the nitrogen loading rate of the synthetic wastewater. The conversion factor *v*N_2_O was calculated with Eq. 3.

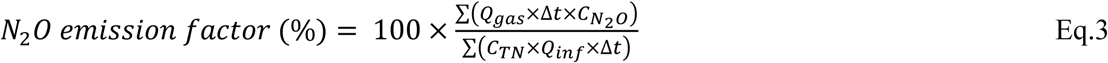

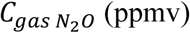= gaseous N_2_O concentration

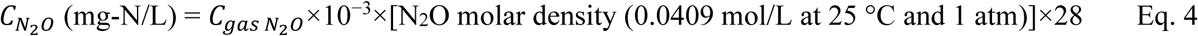

*Q*_*gas*_ (L/min) = the gas flow rate

*C*_*TN*_ (mg-N/L) = total nitrogen concentration of influent

*Q*_*inf*_ (L/min)= the influent flow rate

*Δt* (min) = time interval when the exhausted N_2_O concentration was recorded

### DNA extraction and amplicon sequencing

For microbial analyses, time-series biomass samples were collected from the two MAS reactors (R1 and R2). Total genomic DNA was extracted using a Fast DNA Spin Kit for Soil and a FastPrep-24 instrument (MP-Biomedicals, Santa Ana, CA, USA) following the manufacturer’s protocol. DNA quality was verified with 1% agarose gel electrophoresis and spectrophotometry (NanoDrop 2000, Thermo Fisher Scientific, MA, USA). DNA concentration was determined with the Qubit dsDNA HS Assay Kit (Invitrogen). The V4 region of 16S rRNA gene was amplified using the universal primer set 515F and 806R.^27^ PCR products were cleaned up with an AMPure XP kit (Beckman Coulter, CA), followed by electrophoresis on 0.7% agarose gel and 1.15% Synergel (Diversified Biotech, MA). The DNA gel bands were excised and purified with a Wizard^®^ SV Gel and PCR Clean-Up System (Promega, WI). The final amplicon products were quantified using the Qubit dsDNA HS Assay kit (Invitrogen). Amplicons were pooled in equimolar concentrations and subjected to MiSeq Illumina sequencing using MiSeq Reagent Kit v3 as previously described.^28^

### Microbial community analysis

Sequencing data were generated for 62 samples collected from day 55 to the end of the experiment (day 301) from R1 and R2. Bioinformatic analysis of amplicon sequencing data was performed with QIIME

2 2021.11.^29^ Quality trimming, de-noising, chimera removal, and merging paired-end reads were performed with DADA2 (via q2-dada2).^30^ Sequencing resulted in a table of amplicon sequence variants (ASVs) and their representative sequences. One sample with low sequencing depth was removed from the downstream analysis, leaving a total of 61 samples with sequencing depth greater than 13,750 sequences per sample. Taxonomy of ASVs was assigned using the q2-feature-classifier^31^ classify-sklean naïve Bayes taxonomy classifier against the MiDAS (v4.8.1) database.^32^ The ASV table, taxonomy, and metadata were imported into R (v4.1.2) as a Phyloseq object via the qiime2R package. Alpha-diversity metrics (Shannon and Simpson indices) were calculated with the “estimate_richness” function in the Phyloseq package^33^ at even sequencing depth of the lowest sequences per sample. Principal component analysis (PCA) and redundancy analysis (RDA) were generated using the “amp_ordinate” function of the ampvis2 package^34^ with the default setting. Partial RDA was achieved by the “RDA” function in the vegan package. All significant tests were performed in R. The software PICRUSt2 (v2.4.2) was employed to infer the functional profile of the microbial communities with default setting.^35^ This software places ASVs into a reference tree (20,000 full-length 16S rRNA gene clusters) to infer the gene contents. The gene profile per sample and the mapped pathways were integrated into the final pathway abundances of the sample. All plots were produced with ampvis2 and ggplot2 in R.

### Real-time quantitative PCR (qPCR)

AOMs were quantified by amplifying the functional genes using qPCR. AOB belonging to β-Proteobacteria were quantified with primer set amoA1F-amoA2R^36^ targeting the ammonia monooxygenase α-subunit (*amoA*) gene. Similarly, the primers Arch-amoAF and Arch-amoAR^37^ were used to amplify the *amoA* gene of AOA. Comammox *Nitrospira* clade A, the dominant comammox in wastewater treatment facilities, was quantified using primer set (cmx_amoB-148F and cmx_amoB-485R)^38^ targeting the *amoB* gene. Total bacteria were estimated with the universal primers 1055f and 1392r^39^ targeting the 16S rRNA gene. The details and sequences of all qPCR primers are listed in **Table**

**S1**. A 10-fold serial dilution (10^2^–10^8^ copies/reaction) of plasmid DNAs harboring the targeted genes was prepared to generate the standard curves for absolute quantification. The qPCR was performed with a CFX96 Real-time PCR Detection System (BioRad Laboratories, Hercules, CA, USA). All samples and standards were conducted in triplicate. Detailed PCR conditions were described previously.^40^

### Fluorescence *in situ* hybridization

An activated sludge sample was taken from R1 on day 155 and immediately fixed with paraformaldehyde for 2 h at 4 °C, followed by washing with 1 × PBS and dehydrating by ethanol in series (50%, 80%, and 96% (v/v)). FISH was performed by the protocol described elsewhere.^41^ Oligonucleotide probes were fluorescein isothiocyanate (FITC)-labeled EUBmix^42^ and indocarbocyanine dye Cy3-labeled Nso1225^43^ to detect β-Proteobacteria AOB. The formamide concentration for hybridization was 35%. The probe sequences and formamide concentration can be found at probeBase (https://probebase.csb.univie.ac.at/).

The sample was observed by a confocal laser scanning microscopy (LSM900, Carl Zeiss, Oberkochen, Germany) with an Ar laser (488 nm) and HeNe laser (543 nm).

### Data deposition

Raw 16S rRNA amplicon sequencing data have been deposited to the DNA Data Bank of Japan (DDBJ) Sequence Read Archive (DRA) under accession number DRA014473 (Run: DRR392107-DRR392168; BioSample: SAMD00509971-SAMD00510032; BioProject: PRJDB13859).

## Results and Discussion

### Continuous operation

The average influent TOC and NH_4_^+^ concentrations in the MAS system (R1) were 213.4 ± 79.0 mg-C/L and 361.5 ± 78.5 mg-N/L, respectively. No detectable NO_2_^−^ and NO_3_^−^ were observed in the influent. As shown in **Fig. 1**, organic carbon removal and ammonia retention efficiencies during the entire operation period were 85.5 ± 8.9% and 101.7 ± 24.9%, respectively. Comparable performances (*i*.*e*., 92.1 ± 4.6% for organic carbon removal and 106 ± 39.1% for ammonia retention) were observed in the replicate R2 (**Fig. S2**). During the transition period at the beginning of the experiment, when organic carbon and nitrogen concentrations were fluctuating (**Fig. 1**), NH_4_^+^ retention efficiencies were irregular. Although thorough mass balance examination was required at the beginning, the overall performance of R1 and R2 indicated that the MAS system was successful for organic carbon removal and ammonia retention.

**Figure 1:**
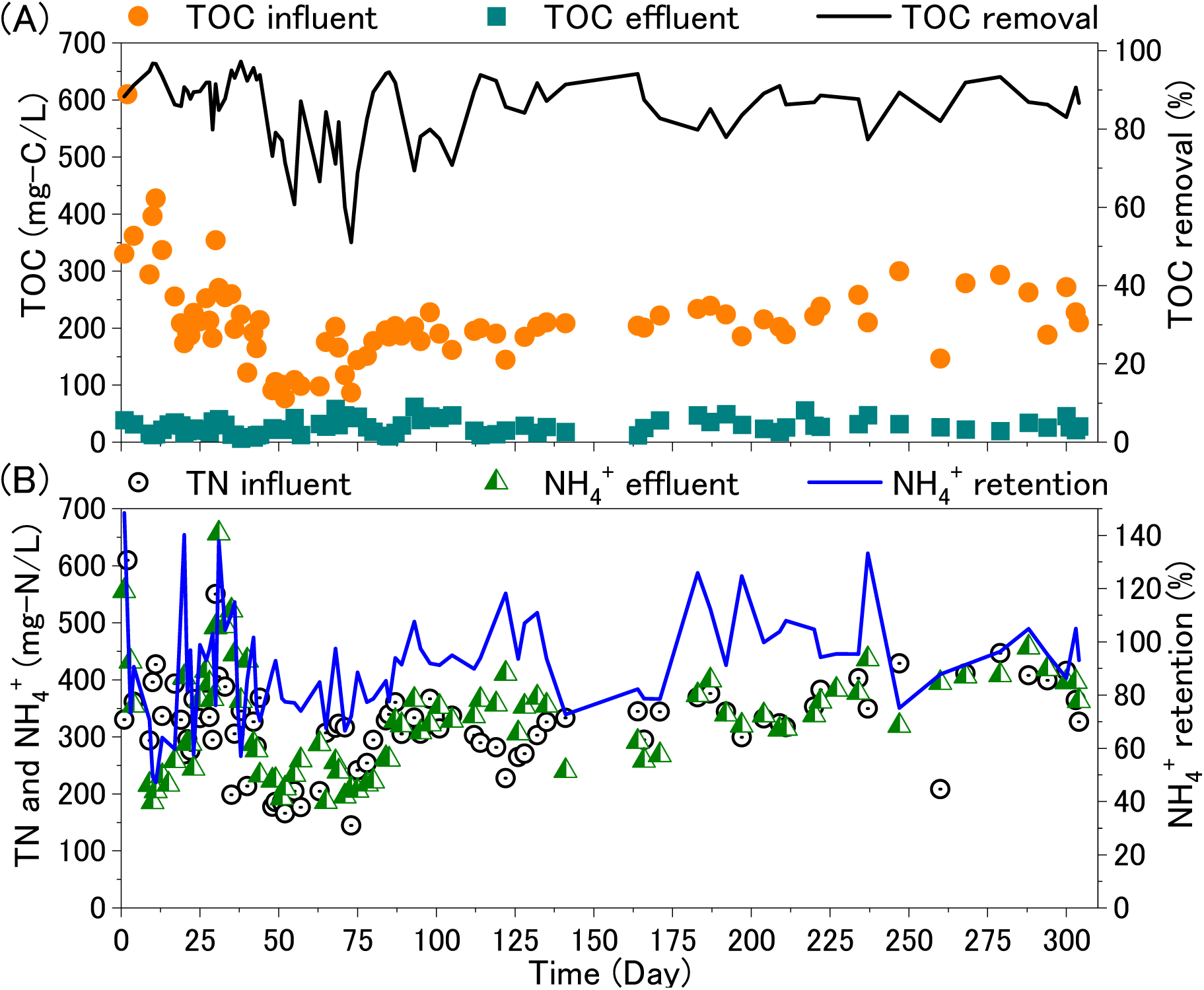
Biological performances by the MAS system (R1): (**A**) TOC concentration and removal efficiency; (**B**) TN/NH_4_^+^ concentration and NH_4_^+^ recovery efficiency.

Although NO_3_^−^ accumulated until day 75, and NO_2_^−^ remained until day 100 (reaching the highest NO_2_^−^ and NO_3_^−^ concentrations of 43.3 mg-N/L and 6.5 mg-N/L, respectively), short SRT (<5 days) and a low DO concentration (setup DO < 0.2 mg/L; actual DO < 0.01 mg/L) successfully suppressed NO_2_^−^ and NO_3_^−^ production. A comparable trend was attained in R2 (**Fig. S3**). The high NO_2_^−^ concentrations were likely caused by the low and fluctuating DO concentrations (**Fig. S4**). Given that AOB have higher affinities for DO than NOB,^16^ suppressed NOB growth was more likely. Previous studies also confirmed this trend.^23, 44^

The experimental conditions likely washed out AOB and NOB after day 125 because the dilution rate (*i*.*e*., the inverse value of the shorter SRT) was shorter than the AOB and NOB-specific growth rates.^16^ A respirometric assay was conducted to determine the ammonia-oxidizing activity of the MAS system during different operating periods. As shown in **Fig. S5**, the ammonia-oxidizing activities in R1 and R2 on day 99 were reduced by 35% and 19% compared with those on day 44. The decreased activity during this period indicated that nitrification was suppressed, congruent with the decreased NO_2_^−^ + NO_3_^−^ and increased NH_4_^+^ in R1 and R2. Subsequently, ammonia-oxidizing activities were scarcely observed on days 168 and 202, consistent with the absence of NO_2_^−^ and NO_3_^−^ in the reactors.

As shown in **Fig. 2**, N_2_O emission factors exceeded 2.0% until day 85 in R1, higher than the default value (1.6%) of the 2019 Refinement to the 2006 IPCC Guidelines.^20^ In contrast, the emission factor sharply decreased after day 130, reaching 0.06%–0.23%. Correlation analysis did not consistently show a positive correlation between the N_2_O emission factor and the accumulated NO_2_^−^ concentrations in the reactors (Pearson correlation of 0.71 [*p*-value = 0.416] for R1 and 0.69 [*p*-value = 0.045] for R2). This inconsistent correlation suggests that nitrite may not be the factor that triggers N_2_O production in the MAS, unlike previous studies of CAS.^45^ Together, the continuous reactor operating conditions (*i*.*e*., SRT < 5 days, DO < 0.2 mg/L, and HRT = 11.2 h) achieved organic carbon removal, ammonia retention, and N_2_O emission mitigation in the MAS system.

**Figure 2:**
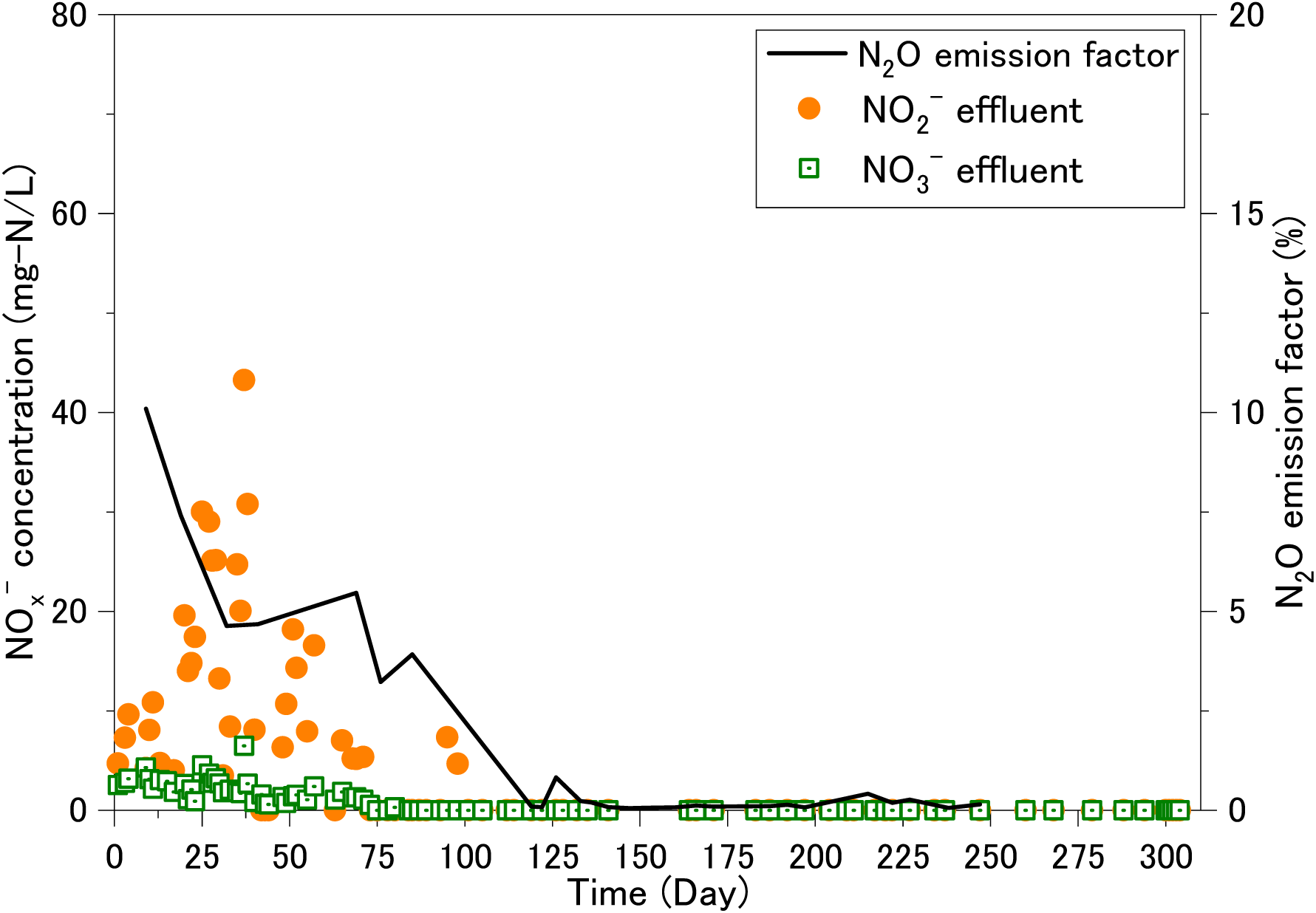
NO_3_^−^ and NO_2_^−^ concentrations in effluent and the N_2_O emission factor by the MAS system (R1).

### N_2_O emission in response to a stepwise change of oxygen loadings

The results of the short-term experiment with stepwise changes in oxygen loading are summarized in **Fig. 3**. In Phase 1, the aeration rate of 2.0 L/min (**Fig. 3A**) triggered ammonia oxidation. The decrease in the aeration rate from 2.0 L/min to 0.5 L/min in Phase 2 resulted in oxygen depletion resulting from organic carbon oxidation and increased NH_4_^+^ concentration (**Fig. 3B**). NO_2_^−^ and NO_3_^−^ concentrations reached 30 mg-N/L and 5 mg-N/L at the end of Phase 1, respectively. During Phase 2, NO_2_^−^ and NO_3_^−^ decreased, NH_4_^+^ increased (**Fig. 3B**), and pH increased while ORP started to decrease (**Fig. 3C**), indicating denitrification. The N_2_O emission factor was 1.4%–2.1% during this transient state (**Fig. 3D**). NO_2_^−^ and NO_3_^−^ were nearly depleted by the end of Phase 2, indicating the endpoint of denitrification. Concurrently, the N_2_O emission factor began to decrease. Anoxic conditions remained during the subsequent 2 h (Phase 3), and NO_2_^−^ and NO_3_^−^ were no longer detected (**Fig. 3B**). The N_2_O emission factor dropped below 0.1% (**Fig. 3D**) and ORP decreased from 100 mV to −150 mV (**Fig. 3C**). Decreased N_2_O emission factor and ORP at the endpoint of denitrification were also reported in a previous study.^46^ Increasing the aeration rate back to 2.0 L/min during Phase 4 resulted in DO concentrations above 1.5 mg/L, initiating ammonia oxidation and decreased NH_4_^+^ concentration (**Fig. 3B**). Subsequently, the ORP and N_2_O emission factor increased to >100 mV and >1.0%, respectively, while pH decreased. ORP and pH profiles can be used to monitor real-time control of nitrification and denitrification,^46^ and this study indicates that the N_2_O emission pattern also reflects the completion of denitrification and the initiation of nitrification. Using the concentration profile of gaseous N_2_O, ORP, and pH is promising for the real-time control of unexpected nitrification, which is one key for ammonia retention in a bioreactor.

**Figure 3:**
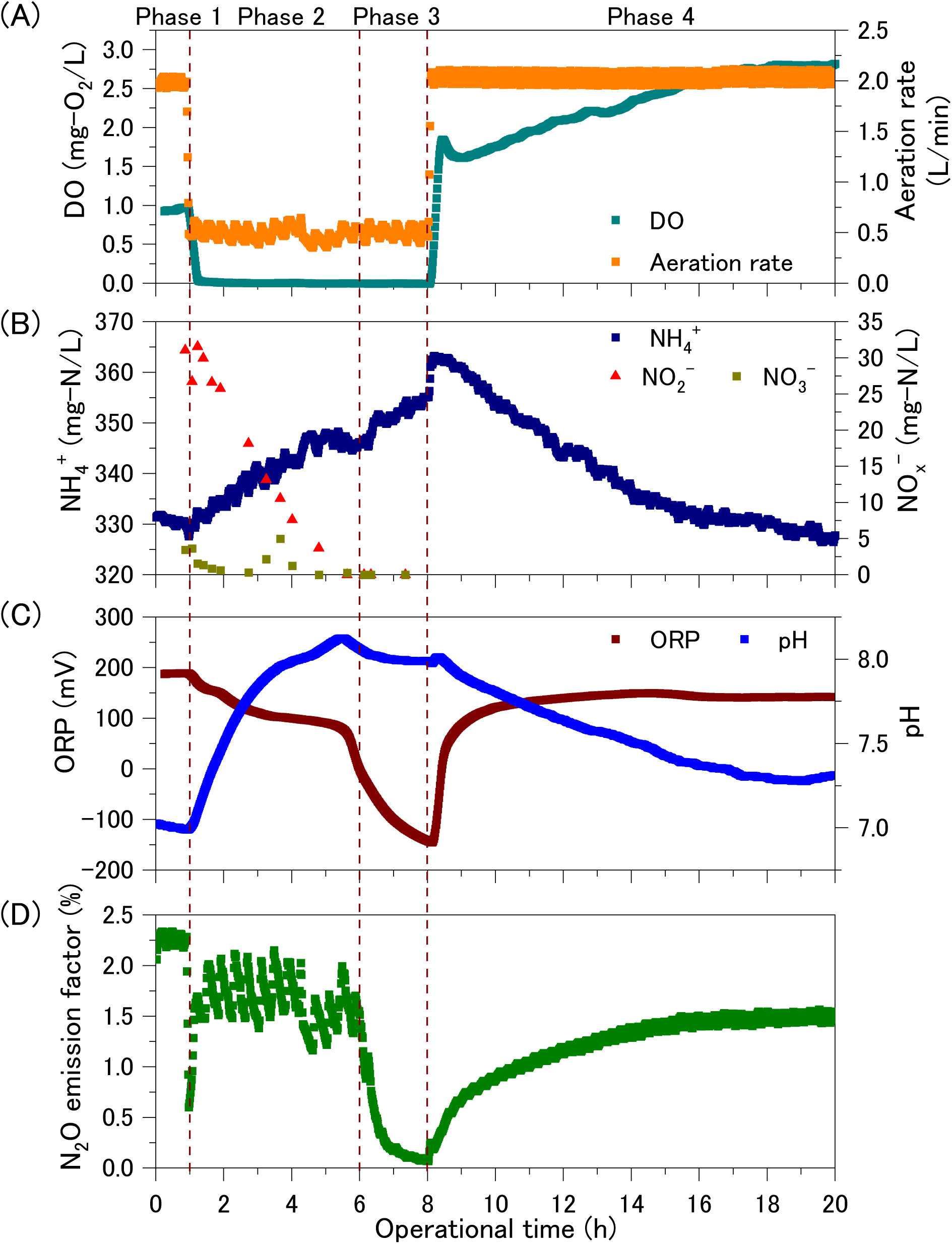
The effects of the stepwise changes in oxygen loading on (A) DO and aeration volume, (B) Profiles of reactive nitrogen compounds, (C) ORP and pH profiles, and (D) N_2_O emission factors. Nitrite and nitrate concentrations were not measured after 8 h.

### Inhibition of ammonia-oxidizing microorganisms in MAS

The application of FISH with Nso1225 and with EUBmix to the microaerophilic activated sludge on day 155 demonstrated that AOB were localized in the deeper zones of microbial flocs (**Fig. S6**). AOB were present as cell aggregates and are often observed in conventional activated sludge (CAS),^47^ indicating that they were not washed out from the system. The growth of heterotrophic bacteria may force AOB into the inner zones of the floc where DO concentration is lower than the bulk liquid. The presence of AOB suggests that the control of oxygen concentration is a key to suppressing ammonia oxidation.

The abundances of AOMs (AOB, comammox *Nitrospira*, and AOA) were tracked by both qPCR and amplicon sequencing. The *amoA* gene copies of β-Proteobacteria AOB were (1.5 × 10^2^)–(1.0 × 10^4^) copies/ng-DNA at the beginning of the experiment, decreased to below the limit of quantification (10^2^ copies/ng-DNA) on day 93, and then remained constant (**Fig. 4**). The change in AOB relative abundance indicated by 16S rRNA gene amplicon sequencing mostly agreed with the qPCR result. The relative abundance of *Nitrosomonadaceae* in β-Proteobacteria AOB was 0.12%–1.49% at the beginning of the experiment (**Fig. 5A**) and then decreased to <0.02% at the end (**Fig. 5A**). In contrast, AOA were not detected throughout the experiment by either qPCR of AOA *amoA* gene (**Fig. 4**) or 16S rRNA gene sequencing. Operating the MAS at low DO concentrations and high organic loading could inhibit the growth of AOB and AOA, which have a lower affinity for oxygen than heterotrophic bacteria;^48^ therefore, AOB and AOA could have been outcompeted for oxygen.

**Figure 4:**
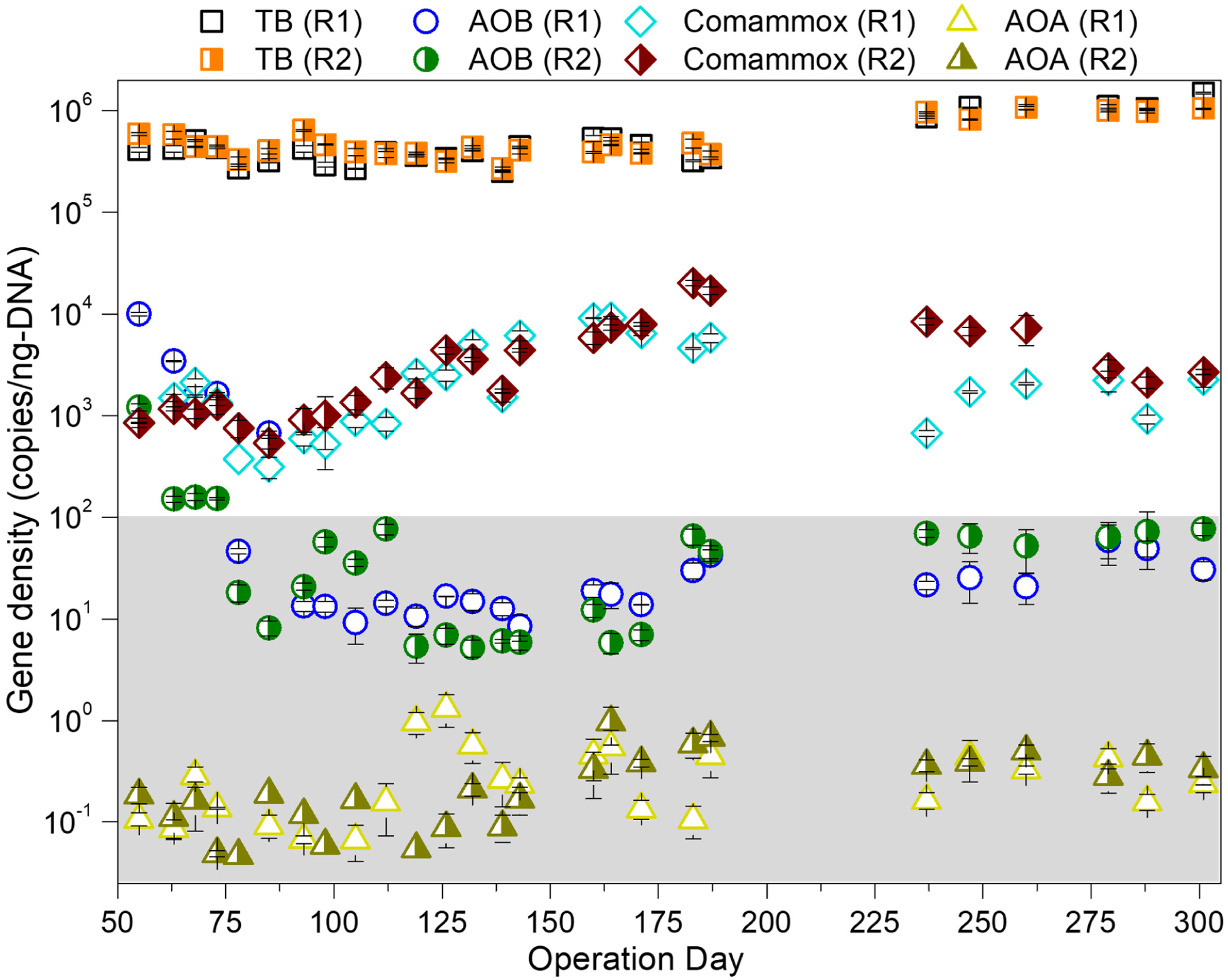
Variations of ammonia monooxygenase gene abundances of AOMs (AOB, AOA, and comammox *Nitrospira*) and total bacteria (TB) in R1 and R2. The area shaded in gray is below the detection limit (10^2^ copies/ng-DNA).

**Figure 5:**
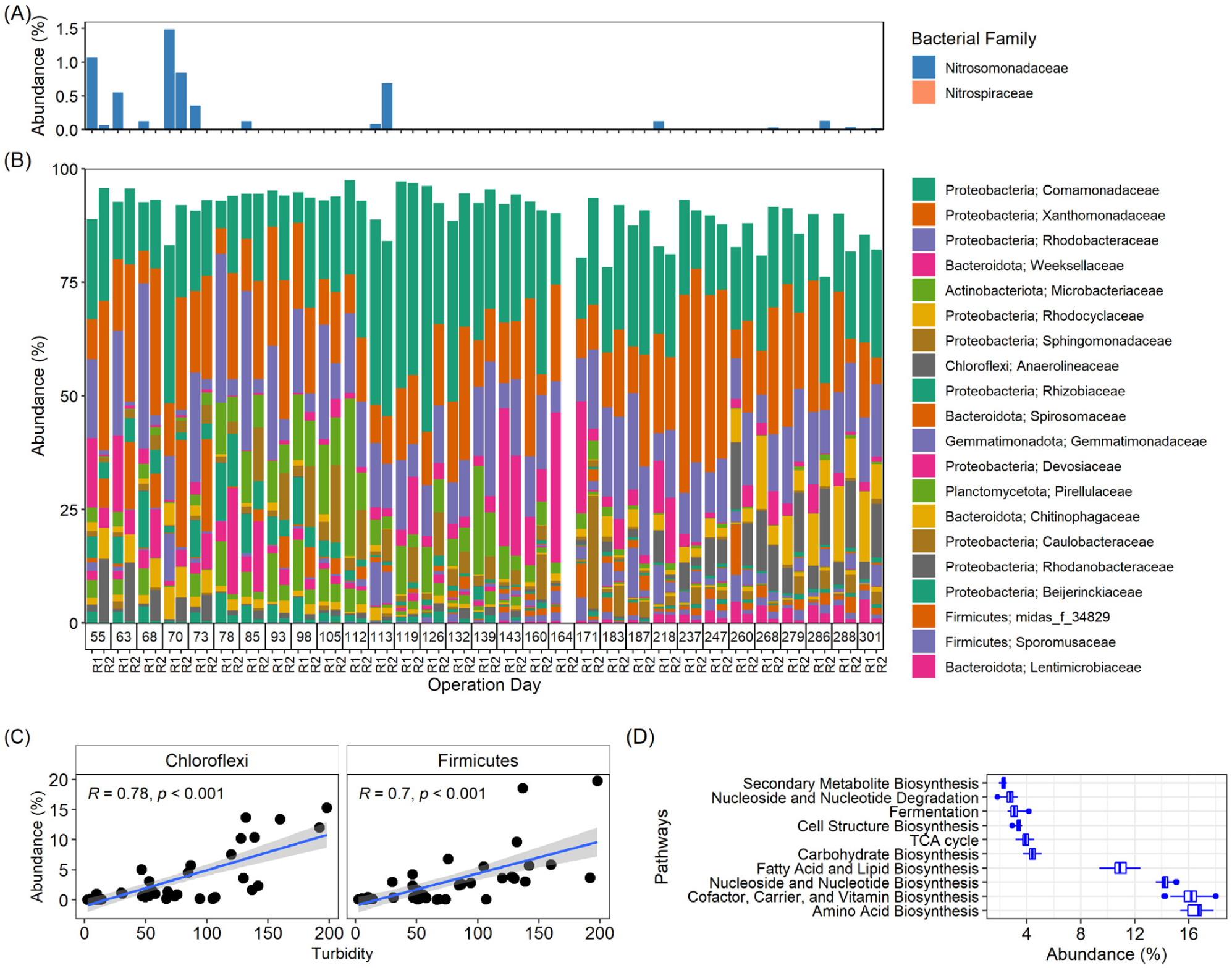
Analysis of microbial community in R1 and R2: (**A**) AOMs detected in the samples; (**B**) The top twenty bacterial families present in the samples; (**C**) Pairwise plots of filamentous bacteria (Firmicutes and Chloroflexi phyla) and turbidity of the bioreactor effluent. (*R* and *p* are Pearson correlation coefficients and significant values, respectively); (**D**) The ten most abundant metabolic pathways in the samples, predicted by PICRUSt2 software. In Panels (**A**) and (**B**), samples of parallel reactors (R1 and R2) are arranged according to the sampling date of operation day (day 55 to day 301). The bacterial families were plotted in the descending order of abundance from top to bottom. On day 164, sample R2 was removed because the number of sequencing reads was below the threshold number (Materials and Methods).

In contrast, the qPCR and amplicon sequencing did not indicate the same population dynamics for *Nitrospira*. The *amoB* gene copies of comammox *Nitrospira* determined by qPCR were (3.1 × 10^2^)–(2.0 × 10^4^) copies/ng-DNA over the whole operational period in R1 and R2 (**Fig. 4**). The presence of the family *Nitrospiraceae*, known to harbor comammox and canonical NOB *Nitrospira*,^49^ was confirmed in only two samples at very low abundances (0.0093% in R2 on day 288; 0.0051% in R1 on day 301) by amplicon sequencing (**Fig. 5A**), which did not agree with the qPCR result. The reason for this discrepancy in *Nitrospira* abundance reported by the two methods is not clear, but the same observation was reported in a recent study.^50^ There is limited knowledge about the diversity and ecophysiology of comammox, and it is not clear if 16S rRNA gene amplicon sequencing or qPCR is more accurate for detecting *Nitrospira* in MAS systems. Of note, the latest report revealed that *Nitrospira* is a core taxon and is universally present in activated sludge,^51^ and was also detected by the V4 region primers used in this study. Primer specificity is an ongoing issue for quantifying comammox *Nitrospira* in the environment.^38,52^ In this study, the operating conditions of the MASs (*i*.*e*., high influent NH_4_^+^ concentration of ∼360 mg-N/L; short SRT of <5 days, except for day 0–132) were not favorable for the growth of comammox *Nitrospira*.^38, 53^ Designing a new primer for the comammox *amoA* gene specifically for the MAS system is crucial. Although further study of the microbial community (*e*.*g*., employing meta-omics methods) is necessary to better understand this issue, both qPCR and 16S rRNA gene amplicon sequencing confirmed the inhibition of AOM growth in the MASs.

### Overall microbial community structure in microaerophilic activated sludge

#### Diversity and microbial assembly

Regarding the α-diversity of all samples, the Shannon and Simpson indices at the lowest sequencing depth (13,750 sequences per sample) were not significantly different between R1 and R2 (Kruskal-Wallis *p*-value > 0.05). While there was a slight variation in Shannon index between time-point samples, the Simpson index was stable during the entire operation period, except for days 237 and 247 (**Fig. S7**). On these days, samples had lower Simpson index than the other samples, indicating a decrease in the evenness of microbial distribution; this observation was verified by examining the microbial composition below.

A Bray−Curtis distance matrix-based PCA was employed to reveal the shift in microbial community structure over time for R1 and R2 (**Fig. S8A**). During the early period of reactor operation (day 55–143), the microbial community structures in R1 and R2 were markedly different, indicated by the distant positions of the R1 and R2 samples. A substantial shift in the microbial community in both bioreactors was initially observed, followed by convergence of the community structure during days 144–218. The samples from days 219–301 grouped closely, indicating similar and stable microbial communities in the R1 and R2 reactors (**Fig. S8A**).

RDA and partial RDA were used to estimate the contribution of operational parameters to the assembly of microbial communities. Seven operational parameters (SRT, turbidity, influent TOC, temperature, ORP, MLSS, and pH) were subjected to permutation tests (999, *p*-value < 0.001) (**Fig. S8B**). All of these factors contributed to 54.8% of the variation in Bray−Curtis dissimilarity between samples (**Table S2**). After controlling for confounding factors, ORP was the most influential parameter (4.7%) in the microbial community variation, followed by temperature (4.5%), influent TOC (4.3%), and SRT (3.1%) (**Table S2**). Previous studies identified temperature, influent organic content, and SRT as the main factors influencing microbial communities in WWTPs;^51, 54, 55^ however, the role of ORP was not directly reported. The dynamic change in the ORP values over a broad range (−340 mV to 220 mV) in this MAS system (**Fig. S4c**) could explain how ORP affects the transitioning microbial communities.

#### Enrichment of a robust organic carbon–consuming microbial community

The twenty most abundant bacterial families in all samples are presented in **Fig. 5B**. Comamonadaceae (22.6 ± 9.5%, n = 61) was the most abundant family, followed by Xanthomonadaceae (18.5 ± 9.9%, n = 61) and Rhodobacteraceae (14.0 ± 8.2%, n = 61). On average, these three bacterial families accounted for more than 50% of the bacterial communities in R1 and R2. The relative abundances were comparable between R1 and R2 (Kruskal−Wallis *p*-value > 0.05, n = 61), suggesting that they play a key role in organic carbon decomposition. These bacterial families are commonly reported as core bacterial phylotypes in activated sludge.^54, 56^ In a comprehensive study of ∼1200 activated sludge samples from 269 WWTPs in 23 countries on six continents,^54^ the abundances of Comamonadaceae members strongly correlated with organic carbon removal rates. Notably, the predominant members of Comamonadaceae in this study and in previous studies^54^ were uncharacterized phylotypes (**Fig. S9A**).

The detected Comamonadaceae consisted of 13 genera, including common members in activated sludge microbiomes (*i*.*e*., *Comamonas, Acidovorax*, and *Simplicispira*) (**Fig. S9A**). The family Xanthomonadaceae in the MAS mainly consisted of two genera, *Thermomonas* and *Luteimonas* (**Fig. S9B**), likely playing an important role in the decomposition of organic carbon. *Thermomonas* was characterized *in situ* as an active denitrifier that can assimilate complex organic carbon sources (*e*.*g*., amino acid and low-molecular-weight nucleic acids) under both aerobic and anaerobic conditions.^57^ A *Luteimonas* isolate obtained from granular sludge in a WWTP was an aerobic heterotroph assimilating different types of carbon sources.^58^ A recent study reported the predominance of *Luteimonas* (>20%) in the effluent of a membrane bioreactor displaying a high organic carbon removal rate.^59^ Xanthomonadaceae had higher relative abundance on days 237–247 (40.6 ± 3.0 %, n = 4). This sharp increase explained the significant decrease in the α-diversity at these timepoints, as discussed earlier (**Fig. S7)**. More specifically, the high abundance of Xanthomonadaceae was caused by the increased abundance of *Thermomonas* (**Fig. S9B**). An increase in temperature may govern the predominance of *Thermomonas* but not of *Luteimonas. Luteimonas* was more abundant in the low-temperature range (<22.5 °C), while *Thermomonas* was more abundant in the high-temperature range (>22.5 °C) (**Fig. S10**).

In this study, Rhodobacteraceae was dominated by an unclassified genus and *Rhodobacter*, followed by 10 other genera (including *Defluviimonas, Paracoccus*, and *Gemmobacter*) (**Fig. S9C**). The Rhodobacteraceae, mainly comprising aerobic heterotrophic non-sulfur purple bacteria, are metabolically versatile and deeply involved in sulfur and carbon biogeochemical cycling.^60^ Cultured *Rhodobacter* species have been used to remove wastewater pollutants. A bioremediation study using high organic load wastewater and the addition of urea under microaerobic conditions showed that a *Rhodobacter* species could intensively biodegrade the organic carbon.^61^ The functions and activities of these taxa in the MAS warrant future investigations.

#### Operation at low DO conditions sustained the growth of filamentous bacteria

A potential downside of the MAS was the poor settling capacity of the activated sludge. The turbidity increased with operational time, as demonstrated by the RDA analysis (**Fig. S8B**). A close examination of the dominant bacteria found that bacterial families in Chloroflexi (Anaerolineaceae) and Firmicutes (midas_f_34829 and *Sporomusaceae*) increased over time (**Fig. 5B**). A pairwise plot revealed strong positive correlations between the relative abundances of Chloroflexi and Firmicutes with effluent turbidity (**Fig. 5C**), which agrees with previous reports.^62^ Many members of these two phyla have been characterized as filamentous bacteria, and their excessive growth can deteriorate the ability of the sludge to settle.^62^ These phyla thrive at low DO, long SRT, low organic loading, and low temperature;^63^ the main factor for excessive growth of filamentous bacteria could be low DO conditions because they have high oxygen affinity and can grow rapidly under oxygen-limited conditions.^64^ Interestingly, the MAS harbored uncharacterized phylotypes of these two phyla, and the uncharacterized genus midas_g_14268 (Anaerolineaceae) in the MiDAS 4 database grew excessively at the end of the experimental period (**Fig. S11**), likely contributing to the poor settleability of the sludge. Close examination and understanding of the ecophysiology of this Chloroflexi phylotype might help mitigate challenges with sludge settling.

#### Metabolic profile of the microbiomes in bioreactors

In total, 51 secondary superclass pathways, classified according to the MetaCyc database,^65^ were predicted for all samples by PICRUSt2. Of these, the ten most abundant pathways accounted for more than 77% (n = 61) of the total relative abundance (**Fig. 5D**). The most abundant pathways were amino acid biosynthesis (16.5 ± 0.7%, n = 61); followed by cofactor, prosthetic group, electron carrier, and vitamin biosynthesis (16.1 ± 0.8%, n = 61); nucleoside and nucleotide biosynthesis (14.3 ± 0.4%, n = 61); and fatty acid and lipid biosynthesis (11.0 ± 0.7%, n = 61). Other abundant pathways included carbohydrate biosynthesis, TCA cycle, cell structure biosynthesis, fermentation, nucleoside and nucleotide degradation, and secondary metabolite biosynthesis (**Fig. 5D**). No pathways involved in ammonia oxidation or denitrification were found, likely because of the long-term exposure to microaerophilic conditions.

#### Implications and future perspectives

An innovative nitrogen recovery strategy requires the development of a simple, robust, and cost-effective technology. Here, we present the proof-of-concept for the MAS system to retain ammonia from high-strength nitrogenous wastewater. The MAS system does not require a specific system configuration but instead retrofits an aeration basin used in the CAS system. A critical challenge for the MAS system is suppressing ammonia oxidation. The long-term and replicate MAS operations demonstrated that operating conditions in the MAS (*i*.*e*., HRT of 11.2 h, SRT of 5 d, setpoint DO concentration of below 0.2 mg/L) allowed high organic carbon removal efficiency and high ammonia retention.

Furthermore, we demonstrated that ORP and gaseous N_2_O concentration from the aeration tank can be used to detect the initiation of nitrification. The molecular microbiological analyses confirmed that the growth of AOMs was suppressed in the MAS, explaining the successful inhibition of ammonia oxidation. In addition, microbial analyses identified the microbial genera responsible for organic carbon removal and poor settleability of activated sludge, which will affect ammonia retention positively and negatively, respectively. Strategies for effective online monitoring of ORP and N_2_O and the managing the microbial communities in the MAS warrant future intensive research. Overall, we demonstrate the feasibility of the MAS as a simple and cost-effective ammonia retention technology. Combining an complementary technology, such as forward osmosis, with the MAS could allow ammonia enrichment and recovery, paving the way for a new nitrogen management paradigm. The development of a tandem system for ammonia retention, enrichment, and recovery is underway.

## Acknowledgments

We would like to acknowledge the Kinu Aqua Station for kindly providing the activated sludge. This paper was based on results obtained through projects of JPNP14004 (Advanced Research Program for Energy and Environmental Technologies) and JPNP18016 commissioned by the New Energy and Industrial Technology Development Organization (NEDO). We thank Ms. Kanako Mori at Tokyo University of Agriculture and Technology for technical assistance in DNA extraction and quantitative PCR, and Ms. Kayoko Suenaga and Mr. Hiroyuki Sakuma at Carl Zeiss Co., Ltd. for technical assistance in a confocal laser scanning microscopy for fluorescence *in situ* hybridization. We thank Tara Penner, MSc, from Edanz (https://jp.edanz.com/ac) for editing a draft of this manuscript.

## Supporting Information

Additional information of materials and methods, Fig. S1–S11, and Table S1–S2 are supplied as Supporting Information.

