## Supplementary data for "Microaerophilic activated sludge system for ammonia recovery from high-strength nitrogenous wastewater: Performance and microbial communities"

### List of Supplementary contents (16 pages in total)

**Fig. S1** Schematic of a laboratory-scale microaerophilic activated sludge system P. S2

**Fig. S2** Replicate reactor performance (R2) P. S3

**Fig. S3** Nitrite and nitrate concentrations of the replicate reactor (R2) P. S4

**Fig. S4** Variations of miscellaneous wastewater constituents in R1 and R2 P. S5

**Fig. S5** Ammonia-oxidizing activity measured by respiration assay P. S6

**Fig. S6** Fluorescence *in situ* hybridization micrograph to visualize the localization of AOB P. S7

**Fig. S7** The Shannon and Simpson indices of microbial communities in R1 and R2 P. S8

**Fig. S8** Principal components and redundancy analyses P. S9

**Fig. S9** Relative abundance at genus level of the three most predominant bacterial families P. S10

**Fig. S10** The abundances of *Thermomonas* and *Luteimonas* as a function of temperature P. S11

**Fig. S11** Classification of phylum Chloroflexi at the genus level P. S12

**Table S1** List of primers used for qPCR P. S13

**Table S2** Fraction of variance of Bray-Curtis dissimilarities P. S14

**References** P. S15

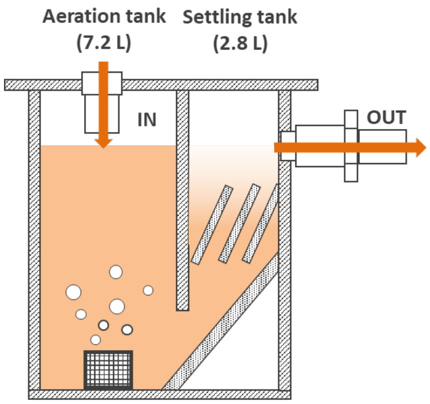


**Figure S1**: Schematic of a laboratory-scale microaerophilic activated sludge system (MAS).

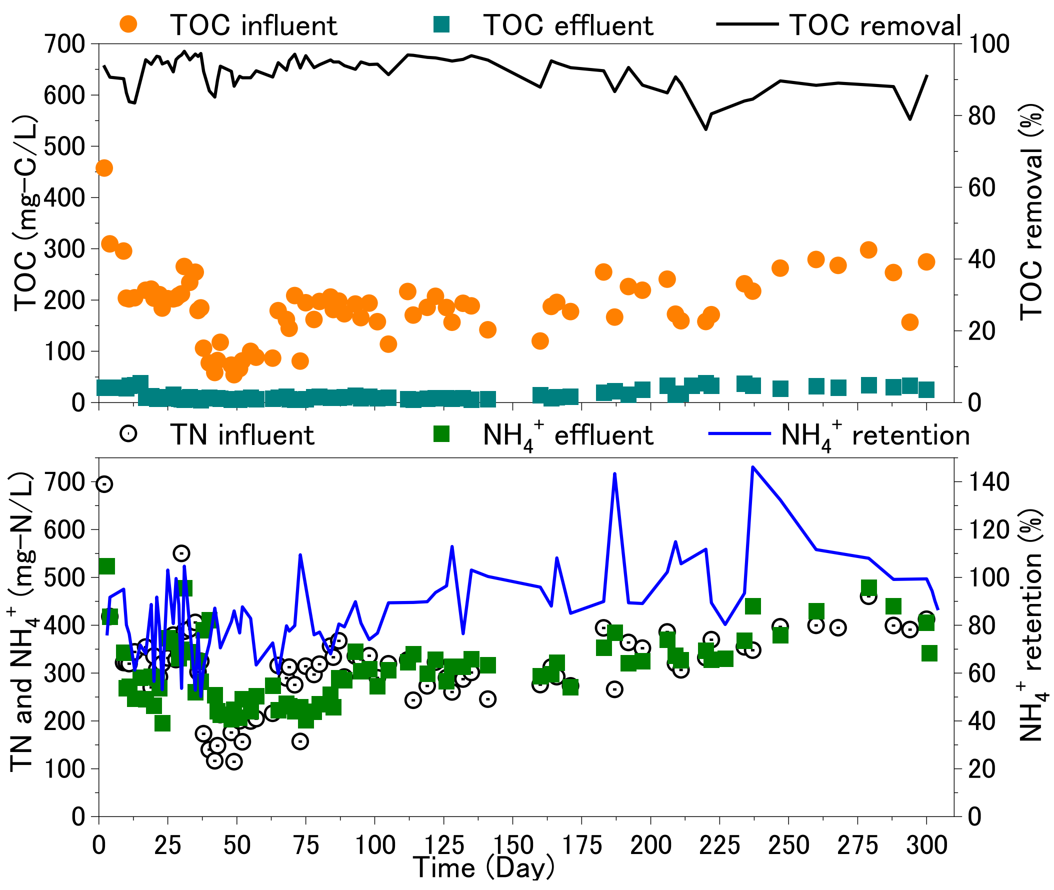


**Figure S2:** Biological performance by Reactor 2 as a replicate: (**A**) TOC concentration and removal efficiency; (**B**) TN and $NH_4^+$ concentrations and $NH_4^+$ recovery efficiency.

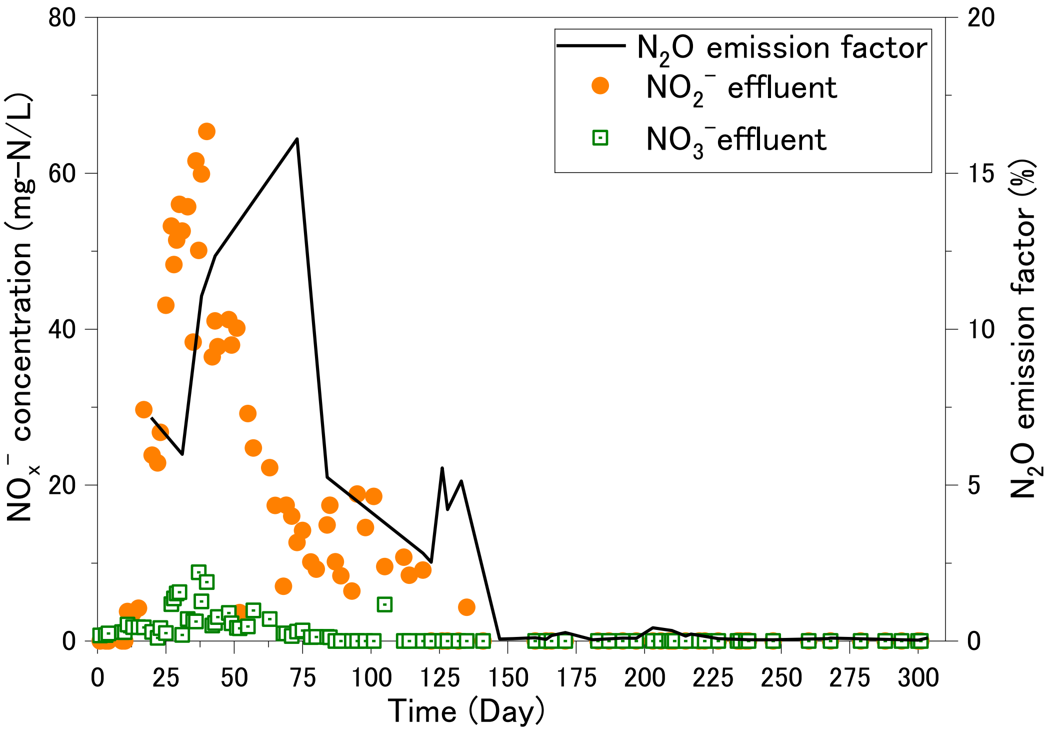


**Figure S3**: $NO_3^-$ and $NO_2^-$ concentrations in effluent and the $N_2O$ emission factor by Reactor 2.

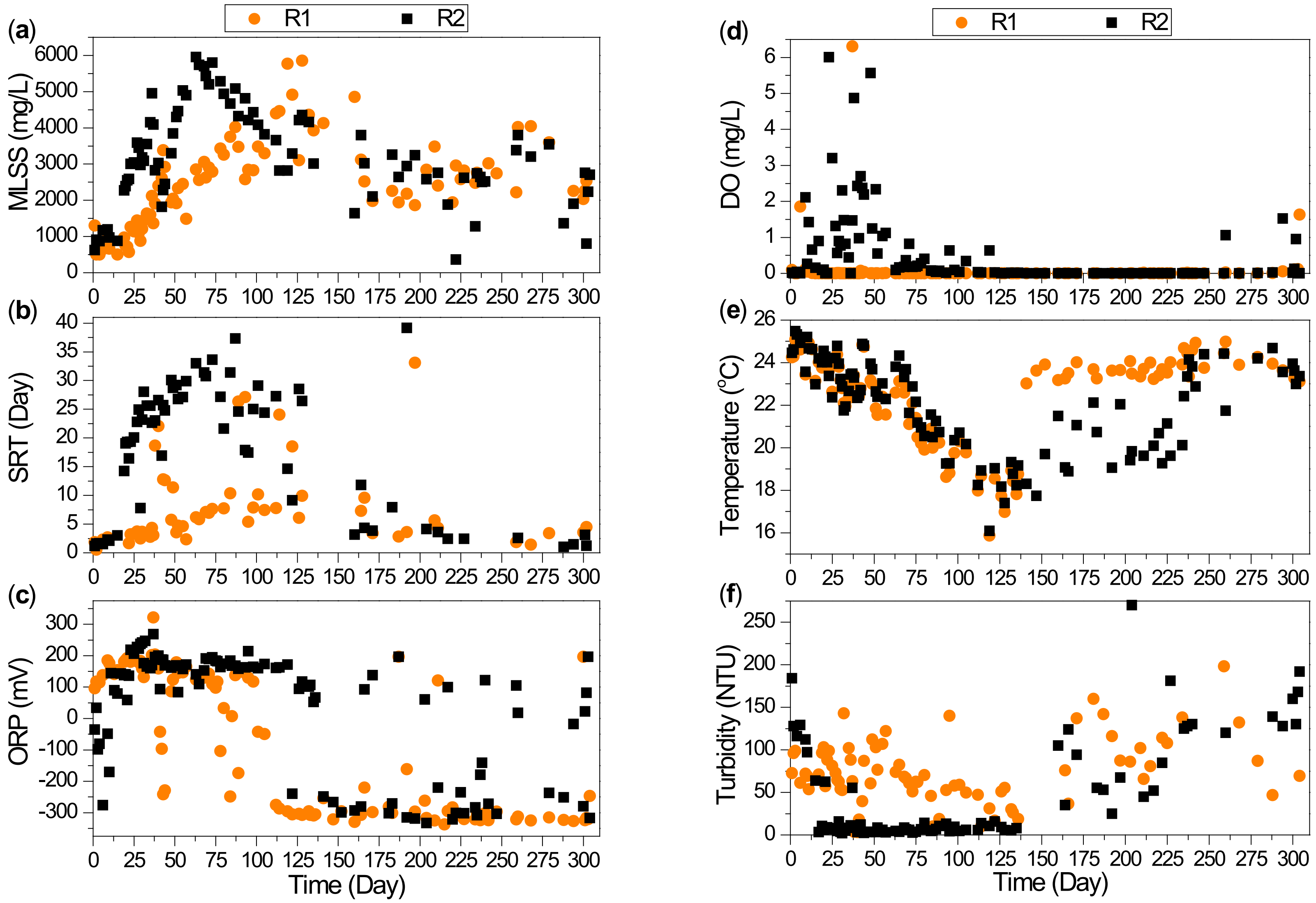


**Figure S4**: Variations of various wastewater constituents in Reactor 1 and Reactor 2.

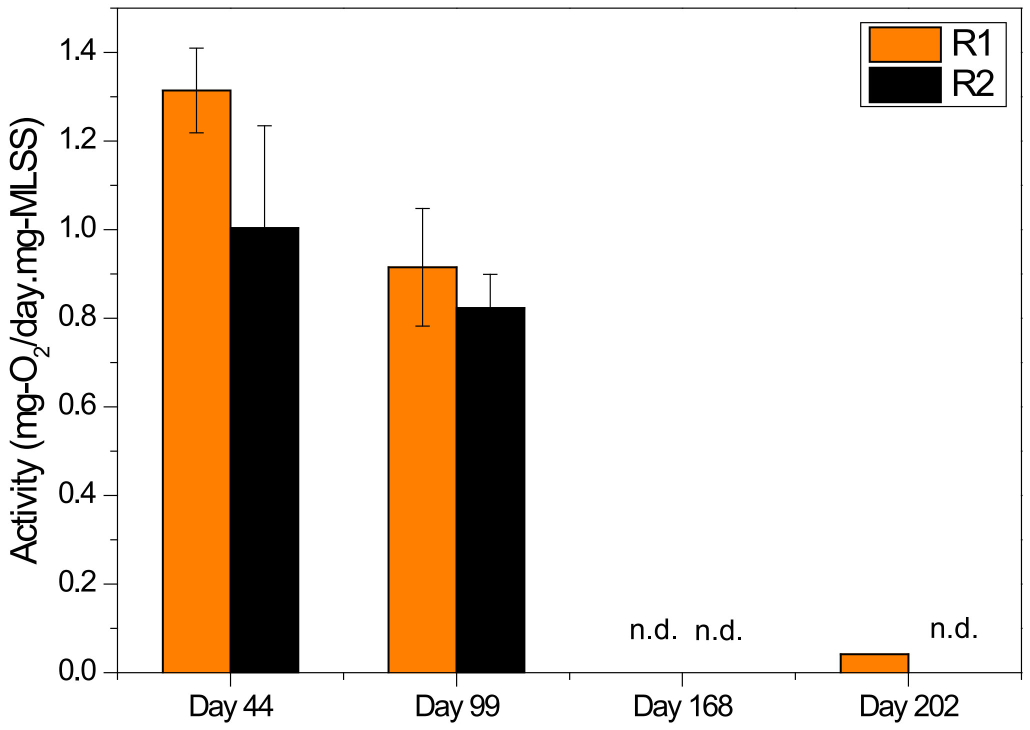


**Figure S5**: Ammonia-oxidizing activity measured by a respiration assay for biomass samples collected on days 44, 99, 168, and 202.

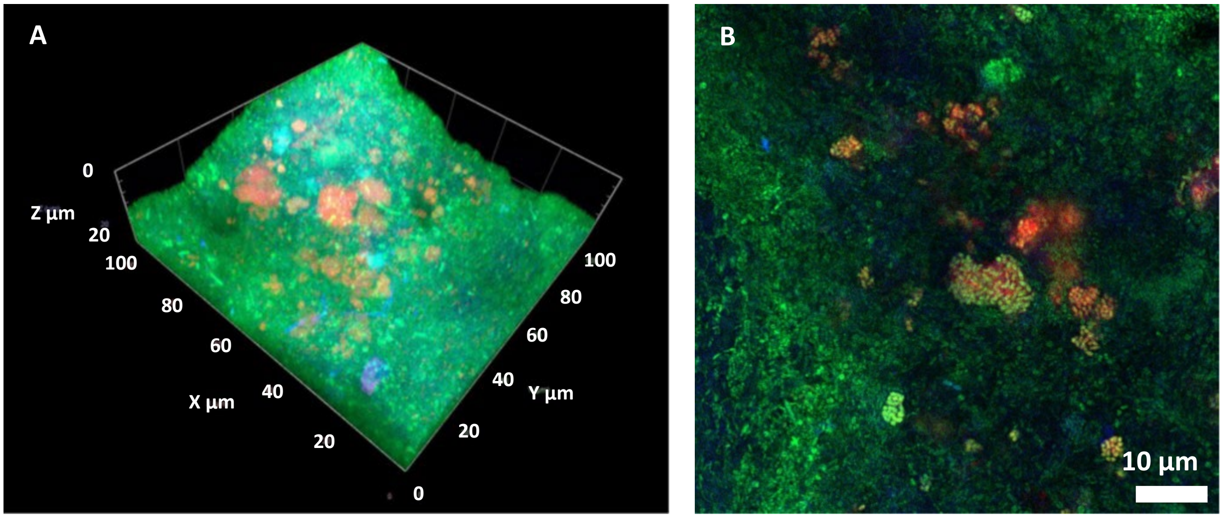


**Figure S6**: FISH micrographs of (A) three-dimensional activated sludge architecture and (B) magnified two-dimensional images visualizing the AOB localization in activated sludge. The sample was hybridized with FITC-labeled EUB mix (Green) and Cy3-labeled Nso1225 (Red) to visualize the β-Proteobacterial AOB (Green + Red = Yellow).

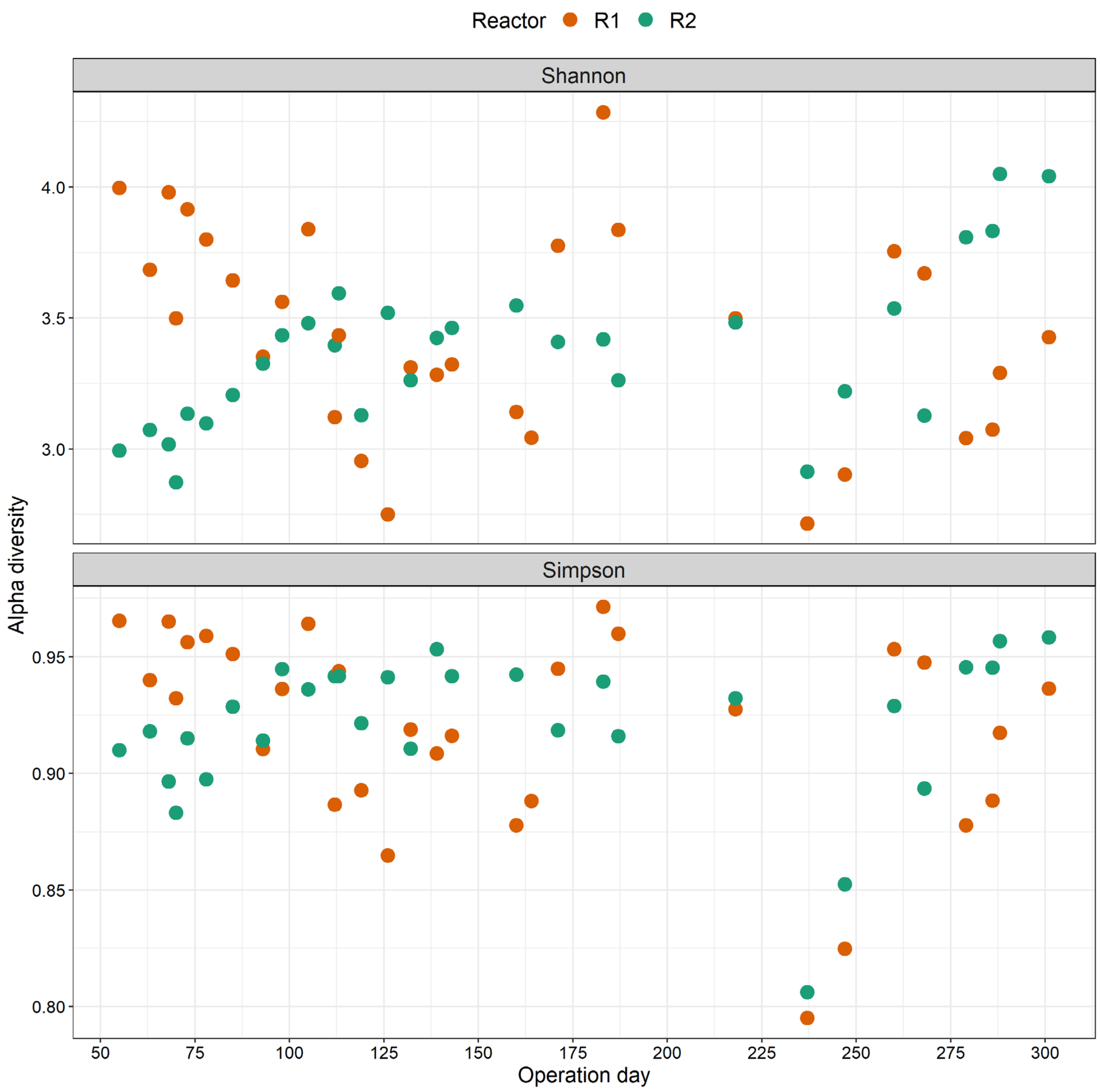


**Figure S7:** α-diversity shown as Shannon and Simpson indices of microbial communities calculated at an even depth of 13750 sequences per sample. The diversity indices were calculated using the "estimate_richness" function (Phyloseq software) by a natural algorithm.

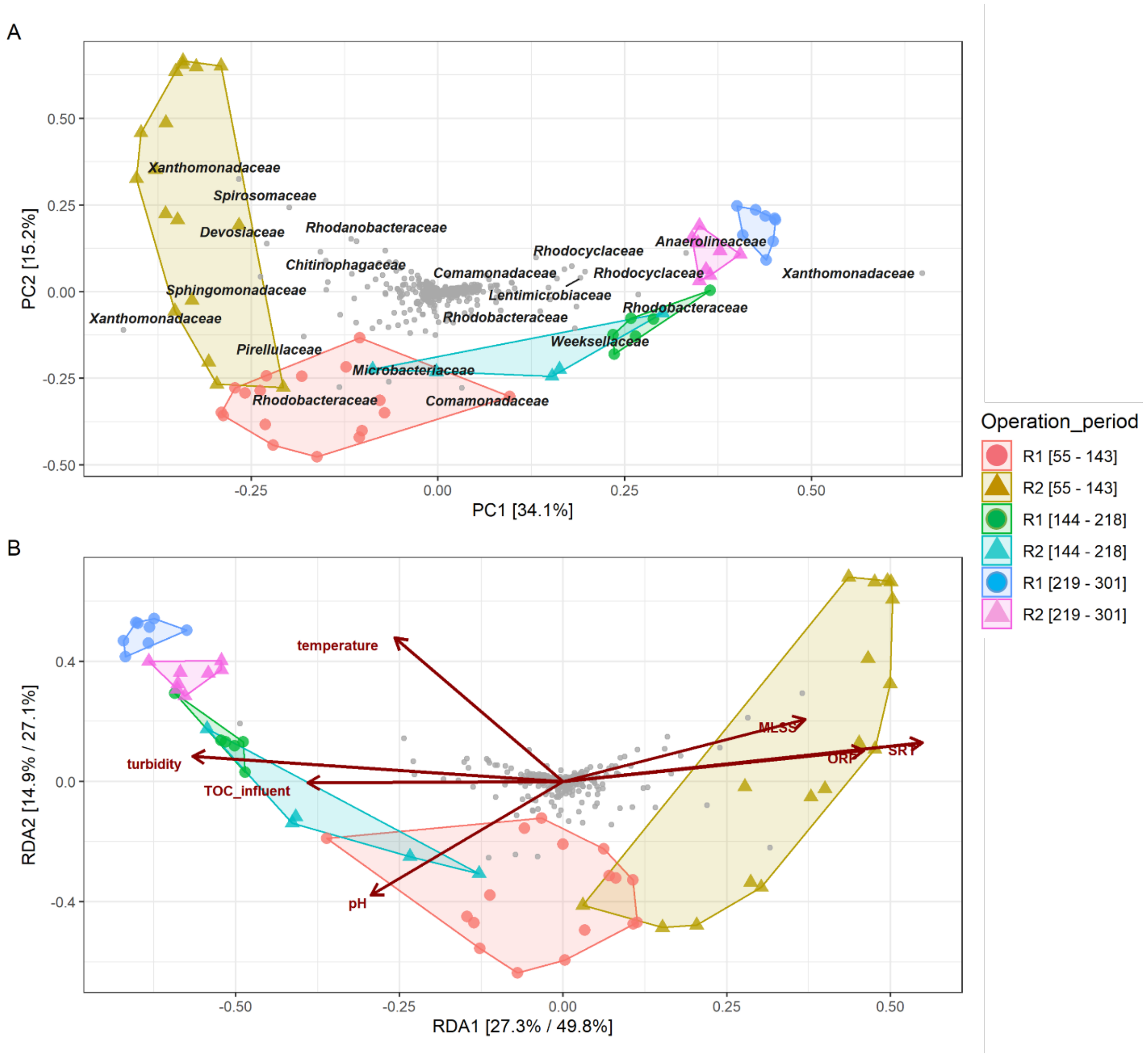


**Figure S8**: (A) Principal components analysis (PCA) and (B) redundancy analysis (RDA) based on Bray-Curtis distance matrices of 61 samples at the single amplicon variant (ASV) level. Prior to the analysis, the ASV table was filtered by removing the ASVs that were not present at more than 0.1% relative abundance in any sample. The data were transformed using a Hellinger method. In panel (A), the relative contribution (eigenvalue) of each axis to the total inertia in the data is indicated in the percent at the axis titles (PCA). The 20 most abundant ASVs were labeled at a family taxonomy level. In panel (B), the relative contribution (eigenvalue) of each axis to the total inertia in the data as well as to the constrained space only, respectively, are indicated in percent at the axis titles (RDA). The length of arrows in the RDA plot were scaled by significance. All the models of operational factors were significant, with *p-values* $< 0.001$ (permutations of 999). Samples were colored according to each reactor's operation period (day), with a circle for R1 and a triangle for R2.

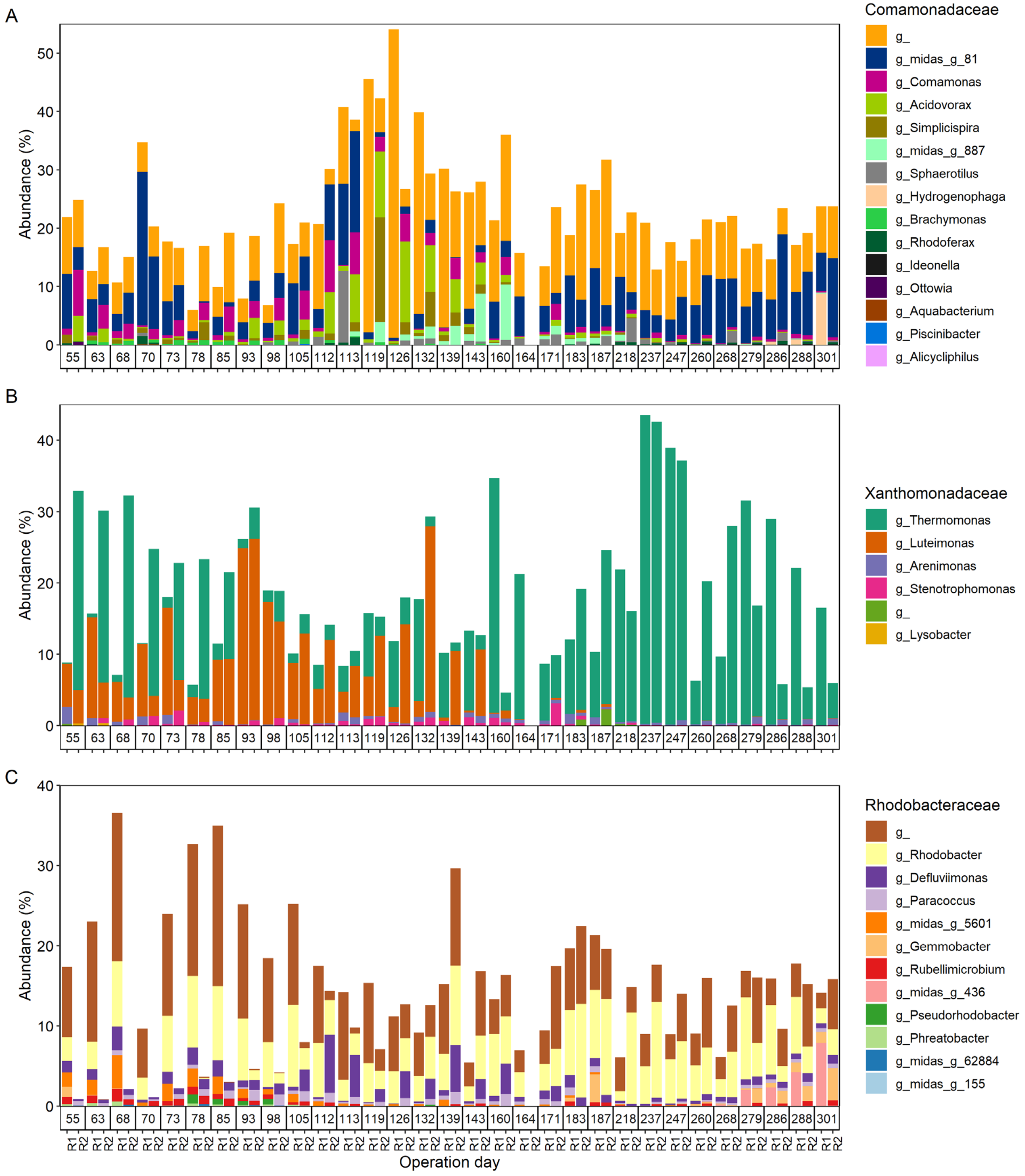


**Figure S9**: Relative abundance at genus level of the three most predominant bacterial families in the microaerophilic activated sludges as presented in **Fig. 5B**.

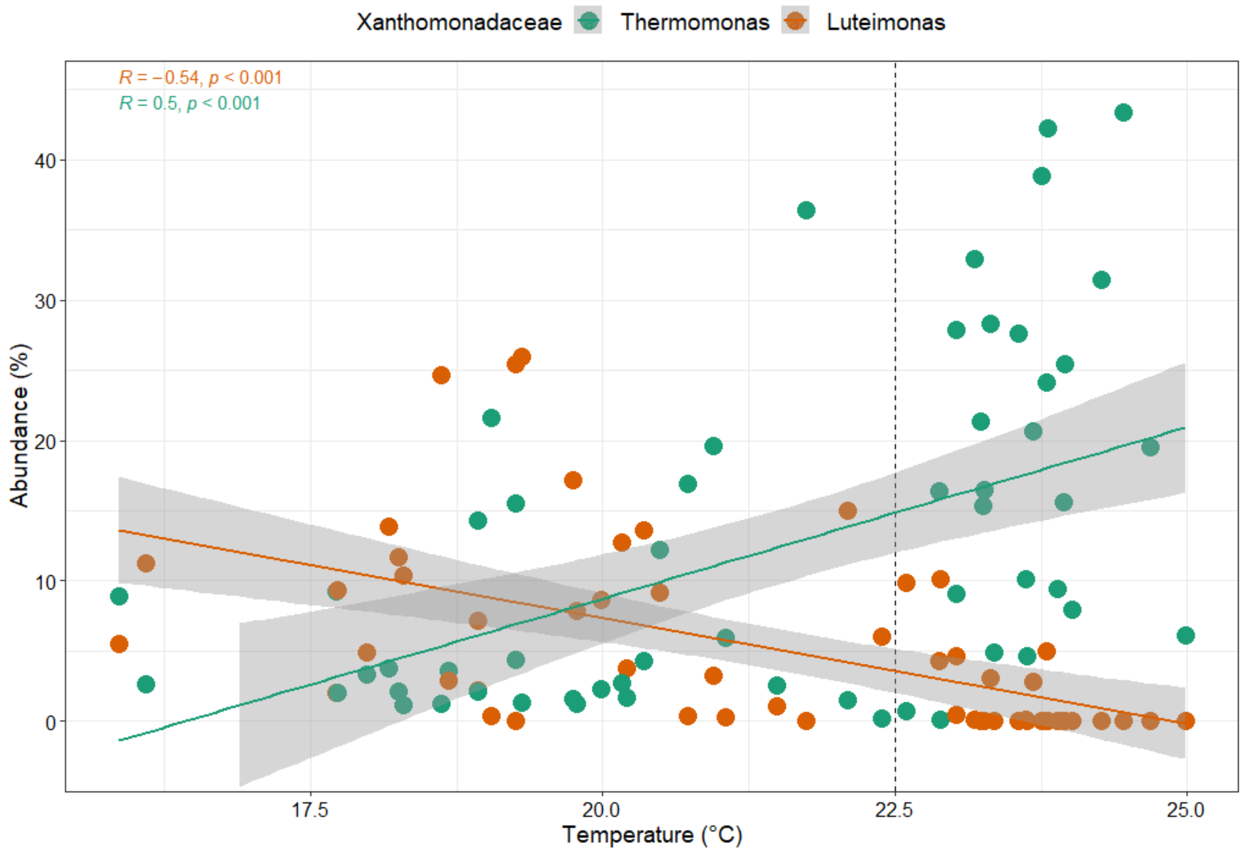


**Figure S10**: The abundances of two dominant genera (*Thermomonas* and *Luteimonas*) belonging to Xanthomonadaceae as a function of temperature. *R* and *p* values indicate Pearson's correlation coefficient and statistical significance, respectively. The grey areas represent the 95% confidence interval for predictions from a linear Pearson's regression.

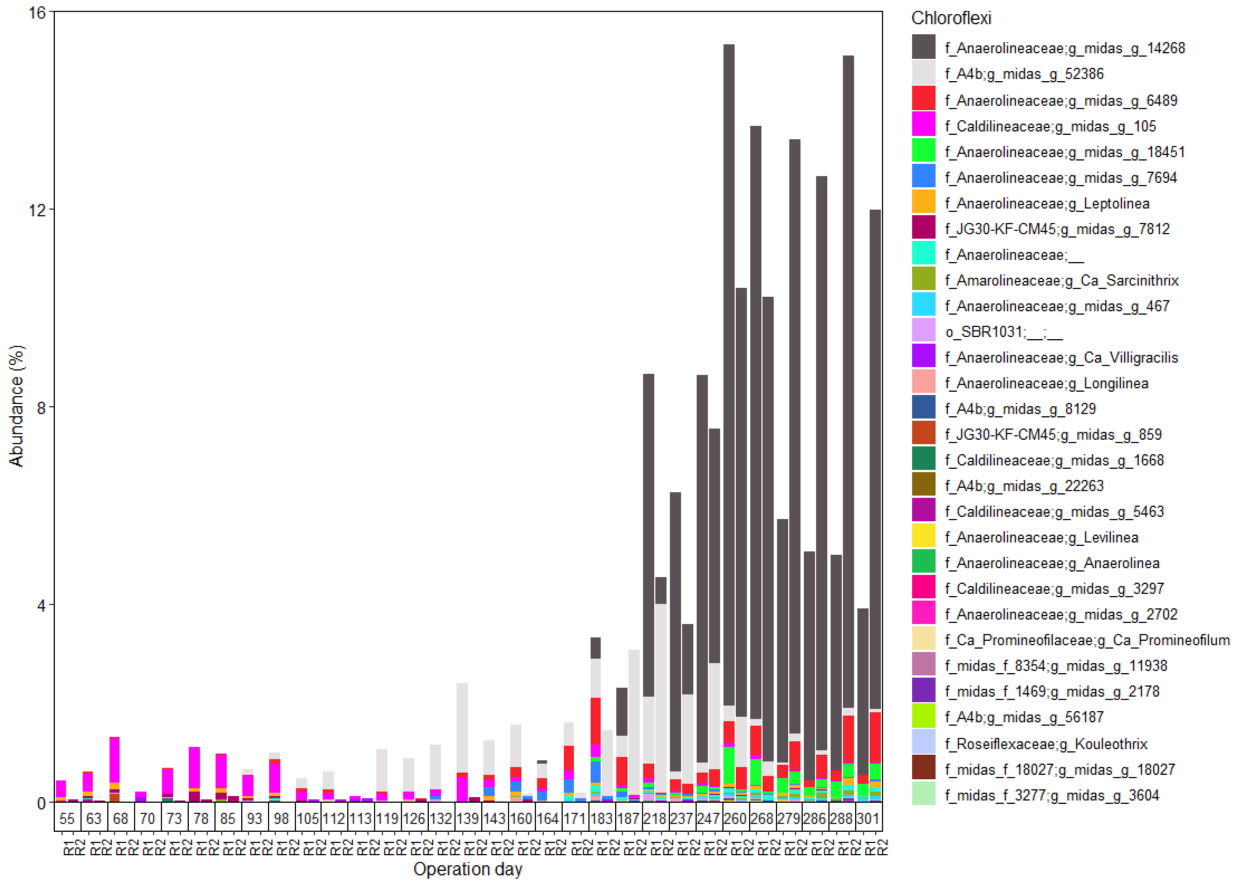


**Figure S11**: Classification of phylum Chloroflexi at the genus level. The genera were plotted in the descending order of abundance from the top to the bottom of the stacked bar.

**Table S1**: Primers used for qPCR

| Primer name | Target | Sequence (5'-3') | Reference |
|---|---|---|---|
| 1055f | Most bacteria | ATG GCT GTC GTC AGC T | Ferris et al. (1996) |
| 1392r | Most bacteria | ACG GGC GGT GTG TAC | Lane et al. (1991) |
| amoA1F | *amoA_Bacteria* | GGGGTTTCTACTGGTGGT | Rotthauwe et al. (1996) |
| amoA2R | *amoA_Bacteria* | CCCCTCKGSAAAGCCTTCTTC | Rotthauwe et al. (1996) |
| cmx_amoB-148F | *amoB_Comammox* | TGGTAYGAYACNGAATGGG | Cotto et al. (2020) |
| cmx_amoB-485R | *amoB_Comammox* | CCCGTGATRTCCATCCA | Cotto et al. (2020) |
| Arch-amoAF | Archaeal *amoA* | STAATGGTCTGGCTTAGACG | Francis et al (2005) |
| Arch-amoAR | Archaeal *amoA* | GCGGCCATCCATCTGTATGT | Francis et al. (2005) |

**Table S2:** Fraction of variance of Bray-Curtis dissimilarities explained by operational parameters. Data were obtained using constrained RDA and partial RDA (for controlling confounding factors). All the models of these operational factors were significant, with a *p-value* < 0.001 after 999 permutations.

| Operational parameters | | SRT | Turbidity | Influent TOC | Temperature | ORP | MLSS | pH | Combination of these factors |
|---|---|---|---|---|---|---|---|---|---|
| Explained variation (%) | Before controlling confounding factors | 23.1 | 22.1 | 18.1 | 17.4 | 15.5 | 13.1 | 8.8 | 54.8 |
| | After controlling confounding factors | 3.1 | 2.1 | 4.3 | 4.5 | 4.7 | 2.4 | 1.5 | |
